# Vector production via mental navigation in the entorhinal cortex

**DOI:** 10.1101/2022.12.15.520640

**Authors:** Sujaya Neupane, Ila Fiete, Mehrdad Jazayeri

## Abstract

A cognitive map is a suitably structured representation that enables an agent to perform novel computations using prior experience, for instance planning a new route in a familiar space^1,2^. Recent work in mammals has found direct evidence for such structured representations in the presence of exogenous sensory inputs in both spatial^3,4^ and non-spatial domains^5–15^. Here, we test a foundational postulate of the original cognitive map theory^1,16^ that cognitive maps are recruited endogenously during mental navigation without external input. We recorded from the entorhinal cortex of monkeys in a mental navigation task that required animals to use a joystick to produce one-dimensional vectors between pairs of visual landmarks without sensory feedback about the intermediate landmarks. Animals’ ability to perform the task and generalize to new pairs indicated that they relied on a structured representation of the landmarks. Task-modulated neurons exhibited periodicity and ramping that matched the temporal structure of the landmarks. Neuron pairs with high periodicity scores had invariant cross-correlation structure, a signature of grid cell continuous attractor states^17– 19^. A basic continuous attractor network model of path integration^20^ augmented with a Hebbian learning mechanism provided an explanation of how the system endogenously recalls landmarks. The model also made an unexpected prediction that endogenous landmarks transiently slow down path integration, reset the dynamics, and thereby, reduce variability. Remarkably, this prediction was borne out of a reanalysis of behavior. Together, our findings connect the structured activity patterns in the entorhinal cortex to the endogenous recruitment of a cognitive map during mental navigation.

## Main text

A hallmark of cognition is the ability to organize experiences into knowledge that can be retrieved flexibly to perform novel mental computations. One way the mammalian brain solves this problem is by establishing cognitive maps that encode spatial, temporal, and other abstract relationships in the environment ^1,2,16,21,22^. The representational building blocks of cognitive maps have been extensively studied in spatial contexts. For example, sensory experiences during both physical and virtual navigation can drive spatially selective responses in the hippocampus and entorhinal cortex ^3,4,23–28^. Non-spatial variables such as temporal relations ^5–9^, value ^10^, social hierarchy ^11^, and abstract stimuli ^12–15^ can also evoke neural responses that reflect the underlying relational structures ^2,29^.

A critical but untested prediction of the cognitive map hypothesis is that the brain can exploit the latent structure of the map in the absence of sensory inputs to perform purely mental computations ^1,16^. To test this idea, we designed a mental navigation task (MNAV) for monkeys in which they used a joystick to move at constant speed along a horizontal line punctuated by six equidistant landmarks, which we refer to as the linear map (Fig 1a). Animals were initially trained to use the joystick to navigate between a subset of possible landmark pairs with all landmarks visible. Subsequently, all landmarks were invisible during movement. As such, animals had to compute and produce displacement vectors on the linear map without sensory feedback.

**Fig. 1:**
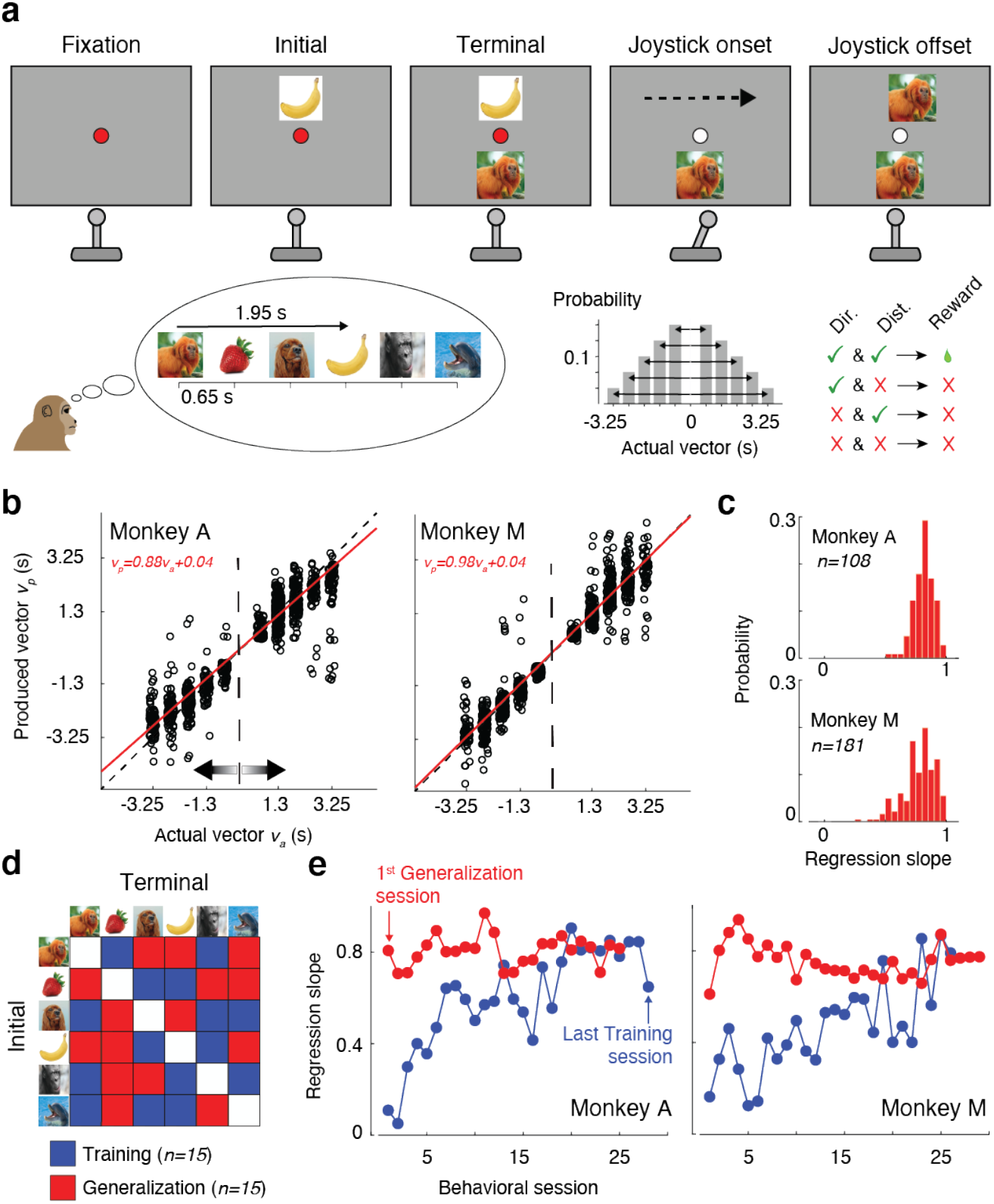
Behavioral task, performance, and generalization. **a**, Mental navigation (MNAV) task. Top: Sequence of events during a trial. Left to right: The animal fixates on a central spot while holding a joystick. The initial landmark is presented above the fixation. The terminal landmark is presented below the fixation. The go cue is presented (change of color of the fixation spot to white). The animal deflects the joystick to the left or right for a duration of time. As soon as the joystick is deflected (onset), the spot above the fixation becomes blank and remains so until the joystick offset. During joystick deflection, landmarks move with a fixed speed (black dashed arrow). After the joystick offset, the landmark closest to the fixation point is revealed. Bottom left: The path consists of six equidistant landmarks with a fixed ordinal position. The animal must learn the sequence and use memory to produce the correct vector by deflecting the joystick for the appropriate length of duration in the appropriate direction. Nearby landmarks are 0.65 s apart. In the example shown, the terminal landmark (image of a tamarin monkey) is three images to the right of the initial landmark (image of banana). Therefore, the animal must deflect the joystick rightward for 1.95 s (3×0.65). Bottom middle: The distribution of directions and distances for randomly chosen initial and terminal landmarks (6×5 conditions). Bottom right: Reward contingency: The animal receives a reward only if both the direction and distance are correct (see Methods for reward window, etc.). **b**, Behavior in a representative session of the two monkeys shown separately. Behavior is measured in terms of the produced vector (*v*_*p*_) as a function of the actual vector (*v*_*a*_) for every trial (open circles). Performance was quantified by the slope of the regression line (red) relating *v*_*p*_ to *v*_*a*_. **c**, Distribution of regression slope across all behavioral sessions. **d**, Initial and terminal landmark pairs used during training (blue) and generalization (red). **e**, Learning trajectory of animals on the training pairs (blue) and on the held-out generalization pairs after the performance on the training pairs had stabilized (regression slope>=0.8, p<.0001 at 95% CI).

We initially familiarized the animals with the task, linear map, and joystick use through a navigate-to-sample task (NTS) (Fig. S1a). On each trial, after fixating a central spot, animals were presented with a target landmark below the fixation point and the linear map above the fixation point. After the fixation point changed color (i.e., ‘Go’ cue), monkeys could deflect the joystick to move the entire linear map horizontally in the corresponding direction. They received a reward for releasing the joystick when the linear map was positioned such that the landmark above the fixation matched the target landmark below (Supplementary Video). Across trials, we randomized the target landmark and the horizontal start position of the sequence while keeping the landmark sequence fixed. While performing NTS, animals gained experience with the relative position of landmarks and the joystick’s speed, the two key variables needed for solving the main MNAV task.

After NTS performance reached criterion (see Methods, Fig. S1b), animals were introduced to the MNAV task (Fig. 1a). MNAV is similar to NTS in that animals use the joystick to move along the linear map to arrive at the target. However, MNAV differs from NTS in that neither before nor after joystick deflection can animals see the entire linear map. First, before joystick deflection, the map is only partially observable via a single landmark right above the fixation point. This modification forces animals to solve the task using their memory of relative landmark positions rather than direct visual input. Second, between joystick onset and offset, all landmarks above the fixation point are invisible, forcing animals to navigate through the sequence using memory. After the joystick offset, the landmark in the sequence closest to the traversed position is revealed, and a reward is provided if this landmark matches the target landmark. If not, the animal is given a second and final chance to make a corrective movement and receive a smaller reward (see Methods). These modifications make MNAV a purely mental navigation task in which animals have to deflect the joystick in the correct direction and for the correct duration to travel between landmarks without any sensory feedback (Supplementary Video). Animals learned to perform MNAV (Fig 1b,c). The produced vectors (*v*_*p*_) quantified in terms of their temporal distance (magnitude) and direction (sign) closely matched the actual vectors (*v*_*a*_).

In principle, MNAV could be solved with two different strategies. One strategy is to treat the task as a stimulus-response memorization problem and learn a lookup table that links each of the 30 (6 permute 2) possible pairs of landmarks to a desired vector. This strategy, which we refer to as model-free, does not require the animals to learn the structured relationships between landmarks. An alternative model-based strategy is to learn and rely on this structure to produce the vectors. The latter strategy involves more sophisticated learning but reduces memory load and offers flexibility when facing new conditions. To evaluate the animals’ strategy, from the outset, we divided the 30 conditions into two appropriately balanced (in terms of direction and distance) disjoint sets, 15 conditions for training the animals, and another 15 to probe their ability to generalize (Fig. 1d). We reasoned that if the animals use a model-free strategy, learning the training set would not confer the knowledge needed to solve the generalization set without additional experience. In contrast, having learned the structure should allow immediate generalization. Evaluating animals’ performance with this paradigm, we found that animals were able to readily generalize with high performance from the very first session (Fig. 1e). This finding suggests that animals solved MNAV using knowledge about the structure of the linear map.

The entorhinal cortex (EC) encodes spatial displacements ^4,23^, responds to spatial and non-spatial variables ^12,14,25,30^, has been hypothesized to support path integration and vector-based navigation ^20,31–33^, and receives sensory input about spatial landmarks ^34^. Accordingly, we hypothesized that EC may play a central role in mental navigation. We recorded spiking activity in EC during MNAV and focused our analysis on the vector production epoch between joystick onset and offset. In both animals, task-modulated neurons (178/614 in monkey A and 194/864 in monkey M) were concentrated in a small region in the posterior EC (Fig. S2). Three prominent features were evident in the activity profile of task-modulated neurons. In some neurons, firing rates were punctuated by transient bumps (Fig. 2a,b). In others, firing rates ramped down and reached different levels depending on distance (Fig. 2a). There were also neurons with a combination of ramping and transient bumps (Fig. 2a). Finally, in many neurons, there was a strong increase in activity before joystick offset. Next, we developed quantitative analyses to examine the relationship between these features and various task and behavioral parameters.

**Fig 2.**
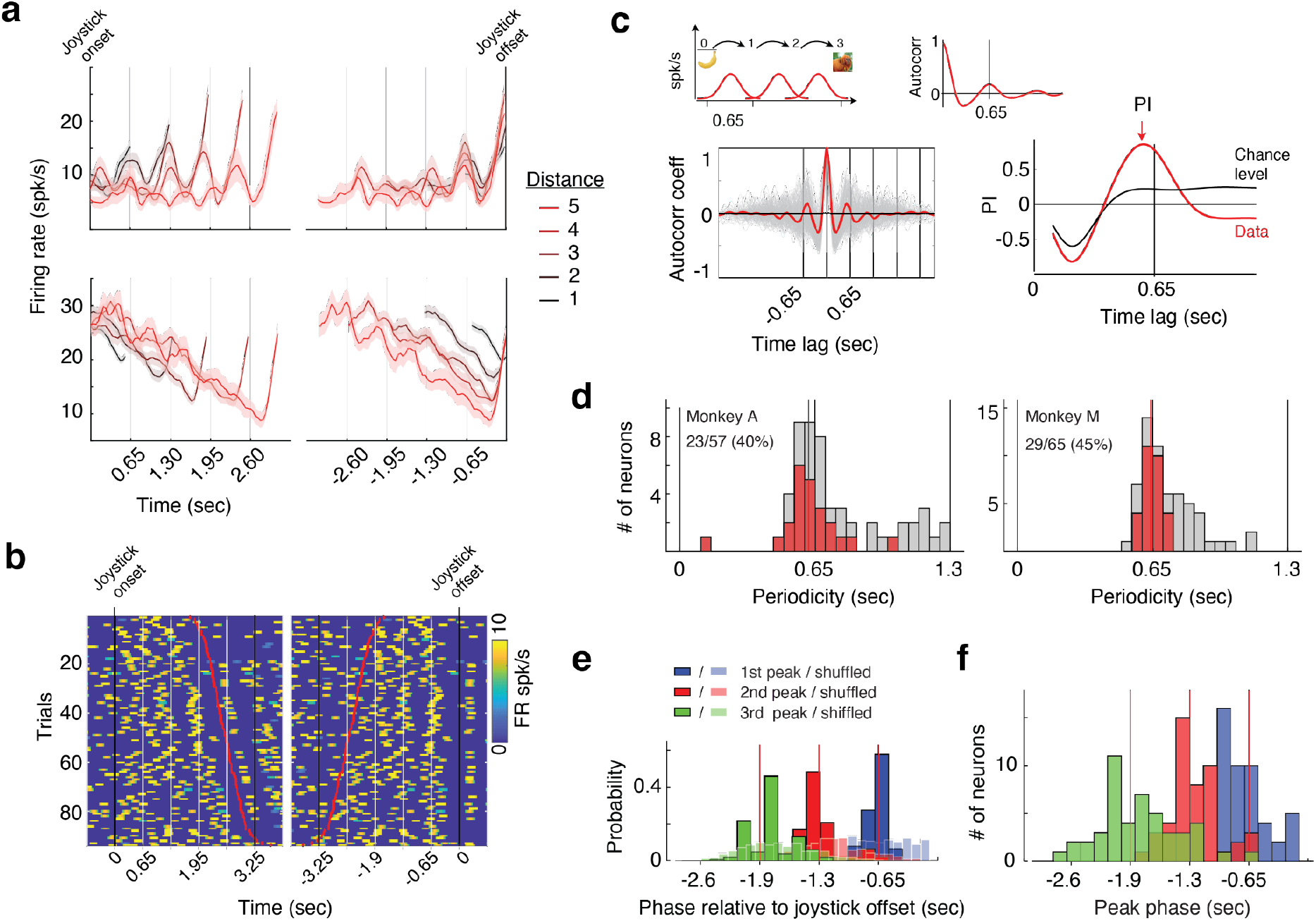
Neural signatures of mental navigation in the entorhinal cortex (EC) **a**, Firing rate of two neurons aligned to joystick onset (left) and offset (right), color-coded by temporal distance. **b**, Raster plot of a neuron for trials with a temporal distance of higher than 3. **c**,(Top) Schematic plot showing how localized activity peaks at landmarks (left) would lead to an autocorrelation function with peaks at multiples of 0.65 sec (right). (Bottom, left) Single-trial (gray) and average (red) autocorrelogram of the neuron displayed in (b). (Bottom, right) Periodicity index (PI) at various time lags (red). Black trace shows the significance threshold set at two standard deviations above chance-level PI derived from surrogate Gaussian Process data (see Methods). **d**, Periodicity values of recorded neurons in one session (red: significant PI, gray: not significant). See Fig S3 for periodicity across all neurons. **e**, Phase distribution of the first (blue), second (red), and third (green) firing rate local maxima before joystick offset. The first peak was estimated within a window of 1000 ms before joystick offset, the second peak was estimated within a window of 1000 ms before the first peak, and the third peak within a window of 1000 ms before the second peak. Phase distributions were calculated based on the mean firing rate bootstrapped over 100 subsamples of trials. The corresponding null distributions were obtained from shuffled spike trains (KS test, p<<.0001). **f**, Distribution of the first (blue), second (red), and third (green) peak phase before joystick offset across neurons.

Considering that landmarks were invisible during navigation and that no other sensory feedback was provided, the presence of transient bumps is striking. We hypothesized that the bumps are internally generated activity modulations associated with the memorized relative position of landmarks. This hypothesis predicts that the time between consecutive bumps should be 0.65 sec, the same as the temporal distance between landmarks. To test this prediction, we computed the autocorrelogram (ACG) of spiking activity for each trial, averaged ACGs across trials, and estimated the time lag associated with the peak of the first side lobe. We considered a neuron to have periodic activity if the value of the ACG at the peak, denoted periodicity index (PI), was two standard deviations above the mean of the null distribution (see Methods, Fig. 2c).

Across sessions, the proportion of neurons with significant PI ranged from 0% to 45% depending on the recording site in EC, with an overall average of 16% (99/614) and 8.5% (74/864) in monkeys A and M, respectively. These percentages are comparable to distance-tuned cells in the medial EC (MEC) in mice running on a treadmill in the dark ^35^ and the percentage of grid cells in candidate regions of rodent MEC ^36–38^ and primates ^25^. Remarkably, across neurons with significant PI, the peak lag was at or near 0.65 sec, which matches the temporal distance between consecutive landmarks (Fig. 2d and Fig. S3a). Complementary analyses revealed no such periodicity in the animals’ eye and hand (joystick) movements (Fig S4), suggesting that the signals were associated with the latent memory of landmarks. This finding provides compelling evidence that EC had a representation of the temporal structure needed for mental navigation between landmarks.

Next, we sought to test the hypothesis that the observed periodicity is linked to behavior. One prediction of this hypothesis is that periodicity should be relatively weak or absent outside the mental navigation epoch. To test this prediction, we compared PI at 0.65 sec between the mental navigation epoch and inter-trial interval (ITI). PI was stronger during the mental navigation epoch (Fig S3b), suggesting that the periodic activity was specific to mental navigation.

Another prediction of this hypothesis is that the trial-by-trial variability in the periodic activity should correspond to variability in behavior. We tested this prediction in two ways. First, we compared PI between correct (rewarded) and incorrect (unrewarded) trials. Doing so, we found that the neurons that were significantly periodic at 0.65 sec had lower PI during error trials (Fig S3c), suggesting that weaker periodicity contributed to committing larger errors. Second, we asked whether the variations of joystick offset time were correlated with phase lag variations of the periodic activity on a trial-by-trial basis. To address this question, we developed an analysis to quantify the phase associated with local activity peaks relative to the time of joystick offset (see Methods). The distribution of the first peak was centered close to 0.65 sec before joystick offset and differed significantly from a null distribution generated by applying random phase shifts to the same spiking data (KS test, p<<.0001) (Fig. 2e). We extended this analysis to earlier times in the navigation epoch and found that the phase distribution for those was centered at near multiples of 0.65 sec (Fig. 2e,f and S5). This finding complements the ACG analysis and further validates the presence of a structured relationship between local peaks and the temporal structure of landmarks. Together, these results establish a tight link between the observed periodicity and behavior and suggest that animals had learned to associate the landmarks with specific phases of the periodic activity.

The other common feature of EC activity during MNAV was a downward ramping activity. To examine the relationship between this ramping activity and task variables, we characterized the degree to which individual neurons had a representation of temporal distances between the initial and terminal landmarks at joystick onset and offset (Fig. 3a). To do so, we performed linear regression to quantify the relationship between temporal distance and firing rates estimated from spiking data within a 100 ms window before joystick onset and offset. Some neurons encoded distance at joystick onset, others at joystick offset, and a small number of neurons had a representation of distance at both onset and offset (Fig. 3b; see Methods).

**Fig 3.**
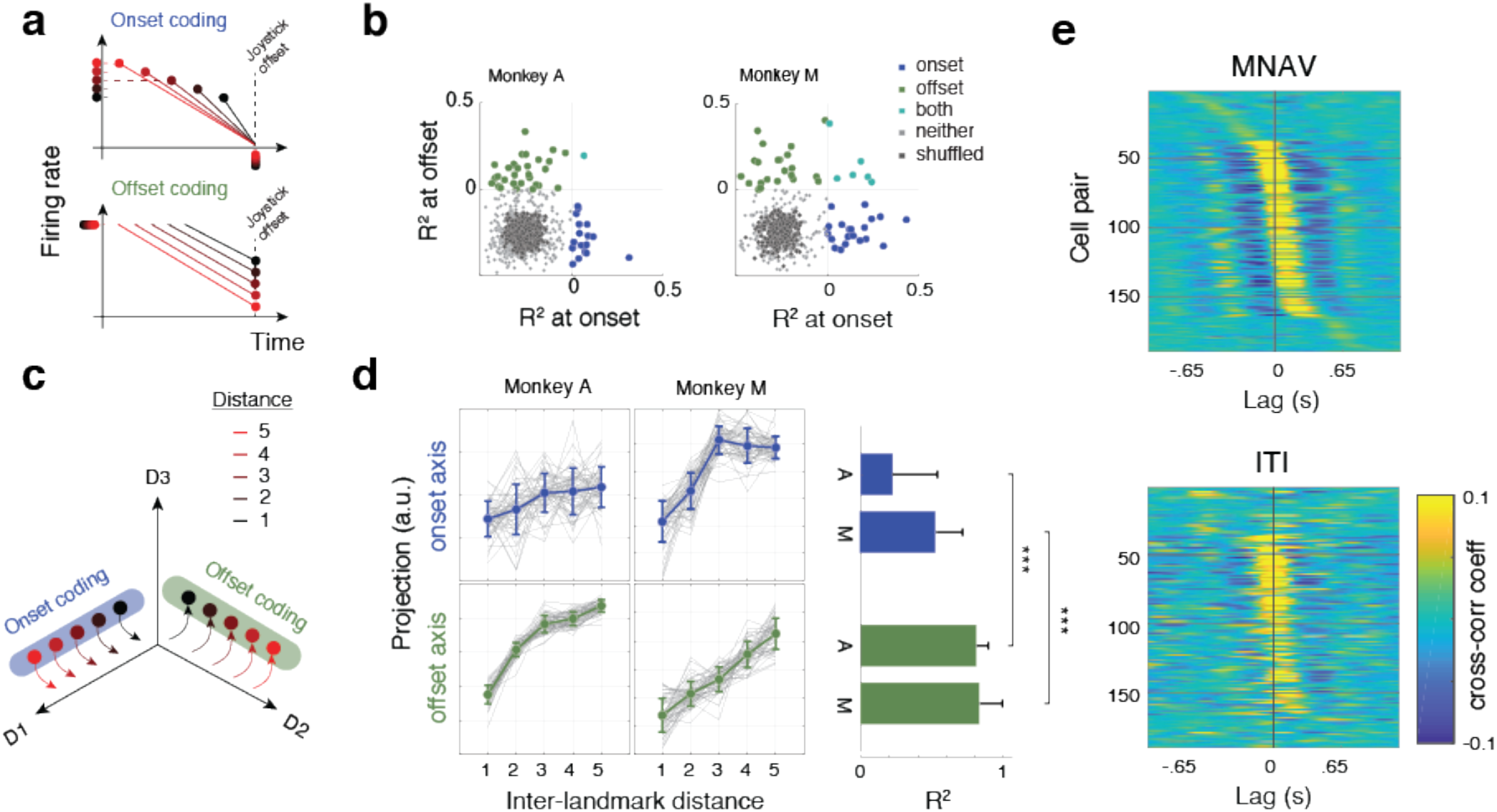
Temporal distance coding and signatures of attractor dynamics in EC. **a**, Schematic depiction of how single neurons might encode temporal distance. To encode temporal distance at onset/offset (onset/offset coding), firing rates should change monotonically with temporal distance at joystick onset/offset. **b**, Scatter plot showing cross-validated explained variance (R^2^) for a linear regression model at onset (abscissa) and offset (ordinate). Different colors represent R^2^ for subsets of neurons with positive temporal distance coding at onset only (blue), at offset only (green), at both onset and offset (turquoise), with no positive coding (light gray), and for a null distribution generated from randomly shuffled regressors (dark gray). **c**, Schematic depiction of how the neural states across the population of neurons might encode temporal distance at onset (blue axis) and/or offset (green axis). **d**, (Left) Projection of held-out neural states onto the axes associated with onset (top, blue) and offset (bottom green) coding estimated using targeted dimensionality reduction. (Right) Goodness-of-fit (R^2^) for onset and offset coding for the two animals (A and M) (two-sided Wilcoxon rank sum test; n=50 bootstraps, Z = −7.21, p<<.0001 for monkey A and n=50 bootstraps, Z=-6.82, p<<.0001 for monkey M). **e**, Cross-correlation structure of 20 simultaneously recorded periodic neurons with the highest periodicity metric (188 pairs) rank ordered based on activity during the navigation epoch (top: MNAV) and plotted with the same order for the inter-trial interval (bottom: ITI) for monkey A (see Fig S6 for monkey M).

Next, we performed complementary targeted-dimensionality reduction analyses ^39^ to quantify onset and offset coding at the population level (Fig. 3c). Similar to the single-neuron analysis, we found cross-validated encoding axes for distance at both joystick onset and offset (Fig. 3d). However, encoding of distance at joystick onset was weak and/or present only for a subset of distances (Fig. 3d, blue). In contrast, population neural states in both animals had a robust linear relationship with distance before joystick offset (Fig. 3d, green). These results reveal two complementary distance codings in EC, one that represents the desired distance before navigation has begun (across initial states), and another that tracks the distance traveled during navigation (across terminal states). Both signals may play a role in navigation. The former allows the system to initialize its dynamics during navigation, and the latter allows the system to track the current state during path integration. While we do not know the source of these signals in EC, we speculate that terminal distance coding may indeed be related to estimating state during path integration as a similar ramping was evident in the position of the joystick during mental navigation (Fig. S4c).

The presence of internally generated periodic activity in task-modulated EC neurons is consistent with velocity integration dynamics in grid cells (GC). Internally generated periodicity is the canonical feature of GC, and these cells acquire this property through attractor dynamics ^17–20,40–42^. Accordingly, we asked whether the observed neural activity in EC is consistent with GC dynamics. A key feature of GC dynamics, besides spatial periodicity, is a broad distribution of relative firing phases across cell pairs that is conserved across conditions, as has been documented experimentally ^17–19^. To test this prediction in our dataset, we measured pair-wise correlations between simultaneously recorded EC neurons with the highest periodicity index in two contexts, during mental navigation and ITI. Consistent with GC dynamics, we found a broad distribution of relative phases between cell pairs and conservation of cell-cell correlations across the MNAV and ITI conditions, across 190 unique cell pairs (Fig. 3e, Fig. S6). Notably, the structure of correlations across the pairs was similar between the two periods (monkey A: r (188) =.84, p<<.0001; monkey M: r (292)=0.86, p<<.0001). These results are consistent with the possibility that the task-modulated EC neurons we recorded are part of the GC system.

These results raise two important questions. First, what circuit mechanisms enable EC to reconstruct the temporal structure of external landmarks internally? Second, what are the implications of this computational strategy for behavior? To address the first question, we adapted a continuous attractor network model of GC ^20^ to model neural dynamics during MNAV. The GC model consists of neurons with lateral interactions that lead to a pattern of repeating bumps. These bumps move in the presence of velocity inputs, thereby supporting path integration. Moreover, inputs associated with external landmarks ^43–48^ can reduce errors accumulated during path integration.

To adapt the GC model to MNAV, we made three assumptions. First, we assumed that either efference copy and/or reafference during joystick deflections provide the velocity input to update the GC state, as has been suggested for path integration in abstract spaces^12^. Second, we assumed that GCs interact with “landmark” neurons (LM) that relay information to GCs about external landmarks when they are present, which is consistent with how landmark information is thought to influence GC activity^34,49^. Finally, we assumed that GC to LM connections are plastic, which is a key component of learning cognitive maps^32,50^. Since landmarks are invisible in MNAV, we reasoned that associative learning at the GC to LM synapses might enable GCs to reconstruct LM activity associated with external landmarks. We tested this idea using a simple model in which an LM unit receives inputs from external landmarks as well as a battery of GC-like input basis sets of different phases and periodicities (i.e., different GC modules) via plastic synapses (Fig. 4a). In the presence of external landmarks, synapses were modified such that GC inputs mimicked the external input, and when the external input was extinguished, the GC drive alone was able to emulate that external drive (Fig. 4b, and Fig. S6). For the specific parameters of the MNAV task (0.65 s between landmarks), the model was able to drive LM with the subset of GCs whose phase and frequency matched the external temporal structure (Fig. 4b). This simple model provides a candidate circuit mechanism for the internal reconstruction of external landmarks in EC.

**Fig 4.**
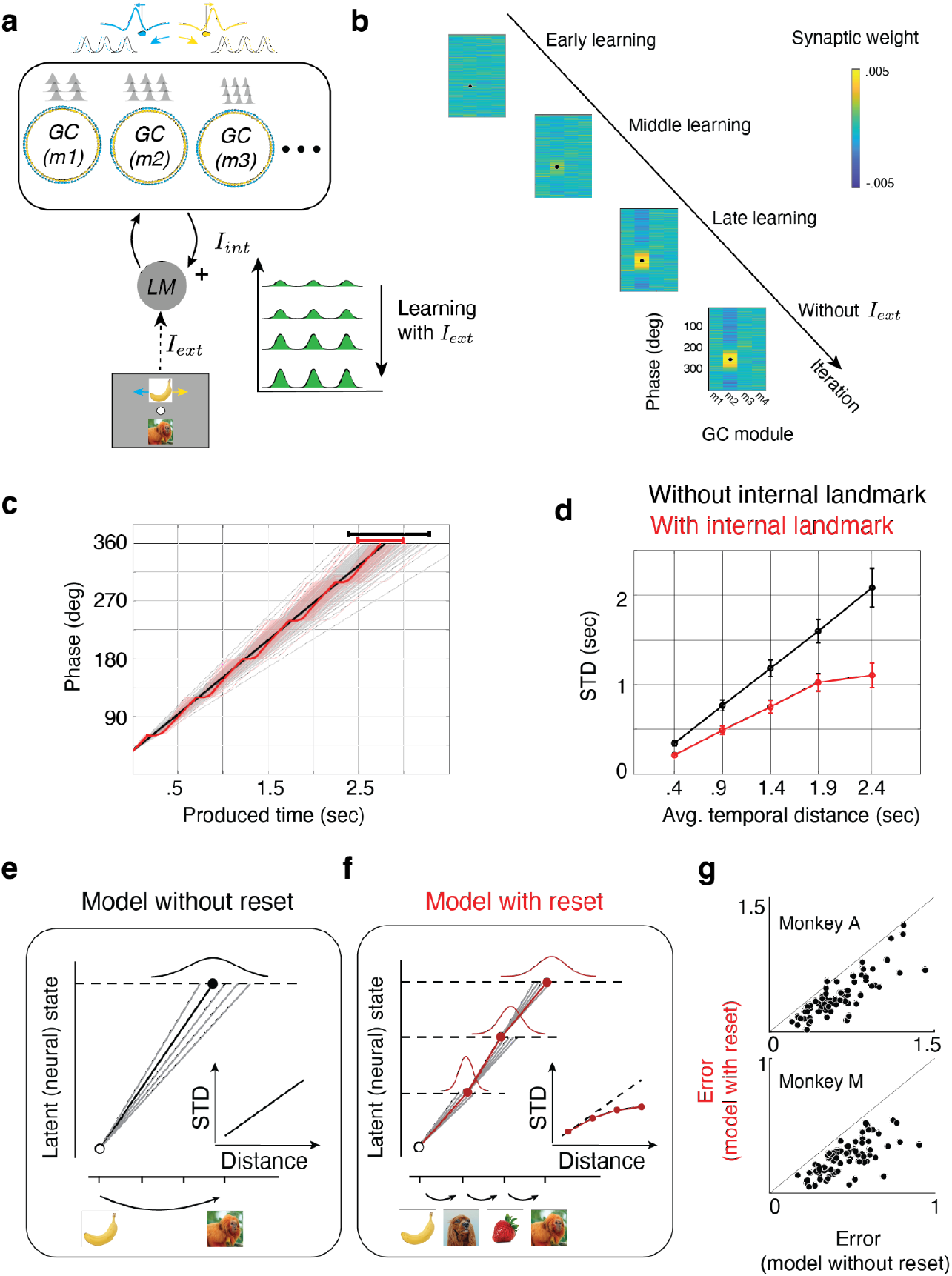
Animals’ behavior is consistent with the dynamics of a continuous attractor network model. **a**, Schematic of a simple continuous attractor network model (box) containing multiple grid-cell (GC) modules (m1, m2, …) with different periodicities and phases (shaded gray inside the box). GC modules receive input from a landmark neuron (LM) that receives both internal input (*I*_*int*_) from GCs and external input (*I*_*ext*_) from visual stimuli. GC neurons have difference-of-Gaussian projection fields whose centers are either shifted clockwise (yellow) or counterclockwise (blue) and receive corresponding velocity inputs (yellow and blue arrows). The synaptic weights from GC to LM undergo Hebbian plasticity (‘+’) and undergo learning both in the presence of *I*_*ext*_ mimicking conditions in NTS and in its absence mimicking conditions in MNAV. The bottom-right panel shows schematically how Hebbian learning selectively strengthens the input from the subset of GCs whose periodicity and phase match that of the external input. **b**, Learning trajectory of GC to LM connections. Synaptic weights from all neurons in all GC modules (m1, m2, ..) are randomized initially (Early learning). In the presence of *I*_*ext*_ with a specific periodicity and phase (black circle), synaptic weights change (Middle and Late learning), strengthening the weights of those GC cells whose periodicity and phase match *I*_*ext*_. The network maintains its selectivity after *I*_*ext*_ is removed (without *I*_*ext*_). **c**, Trajectories of network state across 100 simulations of a 1D CAN model of grid cells with noisy velocity input, with (red) and without (black) landmark inputs. Dark lines show the median trajectory across 100 simulations. **d**, Standard deviation as network state as a function of temporal distance. Standard deviation grows linearly with temporal distance in the absence of internal landmark input (i.e., no reset), and sublinearly in the model with the internal landmarks (i.e., with landmark-induced resets). Inset: Test of sublinearity: Distribution of fitted exponent across 1000 bootstraps (H0: linearity (i.e. exponent=1), one-tailed t-test (999) = −39.32, p<<.0001). **e**,**f**, Two models of behavioral variability. In the model without reset (e, left), the standard deviation increases linearly with distance (e, inset), and in the model with reset (f, left), the standard deviation grows sublinearly (f, inset). Consequently, the overall distribution of produced temporal distances is wider for the no-reset model (top gray Gaussian) than the model with intermediate resets (top red Gaussian). **g**, A Bayesian observer model with the reset (ordinate) provides a better fit to the animal’s behavior compared to a similar model with the reset (abscissa) for nearly all behavioral sessions. Model performance, measured by the difference of mean squared error between animal and model behavior, for two animals (RMSE difference: paired t-test, t(78)=7.89, p<<.0001 for monkey A and t(102)=16.35, p<<.0001 for monkey M)

How do internally generated landmarks influence the dynamics of mental navigation? To address this question, we compared the dynamics of two models, one with the ability to learn and reconstruct landmarks internally, and one without it. The model that did not learn landmarks performed path integration by integrating noisy velocity inputs (Fig. 4c, black) and generated temporal distances whose standard deviation grew linearly with the base interval (Fig. 4d, black), mimicking the well-known scalar property of interval timing^51,52^. In contrast, the dynamics of the model that learned to generate landmarks internally were punctuated with reset-like events coincident with the timing of internally generated landmarks (Fig. 4c, red). In other words, each internal landmark temporarily ‘pinned’ the active bumps and slowed down the movement of the network pattern, acting as a transient reset for the dynamics. This reset mechanism can be readily explained by how LM input interacts with the pattern in the GC network. The LM input stabilizes the formed GC pattern, making it more stable and less responsive to velocity input. The overall effect of the reset induced by the internally generated landmarks was to reduce variability such that the standard deviation of produced temporal distances grew sublinearly with the base interval (Fig. 4d, red). We tested the significance of this sublinearity by fitting the simulation data with a power-law relationship between standard deviation and mean (*std = a*mean*^*b*^ *+ c*; ; H0: b=1; H1: b<1; one-tailed t-test(999) = −39.32, p<<.0001). This observation is consistent with a process of dividing longer intervals into shorter ones to reduce variability^52–54^.

Finally, we returned to the animals’ behavior and asked whether the reduced variability predicted by the presence of intermediate resets provided a better explanation of the variability observed in produced temporal distances. To do so, we constructed two generative Bayesian models (Fig. 4e,f). Both models combined the prior distribution of vector lengths (Fig. 1b) with noisy measurement and used the posterior mean to generate Bayesian estimates (Fig. S7). The two, however, made different assumptions about the form of timing variability during the navigation epoch. The first model assumed that standard deviation scales with the mean, consistent with ignoring landmarks (no reset). The second model assumed that longer temporal distances are divided into multiples of 0.65 s, which leads to a sublinear relationship between the standard deviation and mean (see Methods), consistent with having intermediate resets associated with internal landmarks. Fitting these two models to animals’ behavior, we found that the model with intermediate resets was significantly better than the one without resets at capturing animals’ behavior (Fig. 4g). Together, these results provide a mechanistic understanding of how mental navigation using internal landmarks introduces resets into neural dynamics and how the resulting dynamics reduce behavioral variability.

## Discussion

Our results provide compelling evidence for the recruitment of a cognitive map in EC during mental navigation^1^. EC neurons generated endogenous periodic activity that matched the temporal organization of memorized landmarks. This periodic activity is reminiscent of grid cell activity in rodents^17–19,55^ and may arise from functionally homologous continuous attractor architectures. However, there are notable differences that deserve further scrutiny. First, the velocity input in our experiment is internally generated. This creates a need for the system to calibrate the velocity gain or change the attractor landscape or both so that traversing the landscape would generate the appropriate periodicity. We verified the plausibility of this scheme with Hebbian learning, but other synaptic plasticity mechanisms found in this system might be better candidates^56,57^. Second, many EC neurons exhibited ramping activity, which is not observed in EC neurons that also exhibit periodic activity. This ramping activity may come from two sources. First, the task-modulated EC neurons in our experiment may be among the cells with heterogeneous tuning that inherit their properties through recurrent interactions with other neurons^58^ in different parts of EC^7–9,28,59–62^ and hippocampus^5,6^. Alternatively, the ramping might reflect input from other brain areas that encode upcoming events via ramping^63–68^. The presence of multiple timing mechanisms may be important for calibration and learning^69^.

Our modeling work also has implications and makes predictions for future experiments. First, plastic connectivity between EC and putative ‘landmark’ cells is a salient computational assumption of several recent models of hippocampus and EC^32,43–47,50^. However, its biological basis remains untested. Second, a prediction of our model is the network should be able to accommodate new joystick speeds if it could readily adjust the gain of the internal velocity input. This is not possible in our current experiment but modifying the task slightly to provide visual feedback about the speed (e.g., by a flow field) could be used to test this prediction. Third, the learning component of our model predicts that the same machinery and mechanisms can support mental navigation in the presence of irregular landmarks. The only predicted difference is that internal reconstruction of the arrangement of the landmarks would require a weighted sum of multiple grid modules.

Finally, we speculate about the implications of our findings for other behavioral contexts and neural systems. Many behaviors, such as silent counting and mental rehearsal, involve traversing through structured memories without sensory input. While we know nearly nothing about the precise neural mechanisms of these high-level cognitive behaviors, we note that they have computationally analogous components to our mental navigation task. For example, silent counting may rely on dynamics similar to what we have discovered in EC: a timer with intermittent resets. If so, the behavior may rely on a continuous attractor neural system that treats silent counts as abstract landmarks. This would also explain why counting reduces variability^52^. With these considerations in mind, we hope our work will contribute to circuit-level understanding of cognitive processes within the memory system.

## Methods

All experimental procedures conformed to the guidelines of the National Institutes of Health and were approved by the Committee of Animal Care at the Massachusetts Institute of Technology. Experiments involved two male, awake, behaving monkeys (species: M. mulatta; ID: A and M; weight: 8.4kg and 11.5kg; age: 6 and 11 years old). Animals were head-restrained and seated comfortably in a dark and quiet room, and viewed stimuli on a 23-inch monitor (refresh rate: 60 Hz). Eye movements were registered by an infrared camera and sampled at 1kHz (Eyelink 1000, SR Research Ltd, Ontario, Canada). Hand movements were registered by a custom single-axis potentiometer-controlled joystick whose voltage output was sampled at 1kHz (PCIe6251 board, National Instruments, TX). The MWorks software package (https://mworks-project.org) was used to present stimuli and to register hand and eye position. We used 32- and 64-channel laminar probes (V-probe, Plexon Inc., TX) for neurophysiology recordings driven by a motorized micromanipulator (Narasighe Inc.) through a bio-compatible cranial implant. Analysis of both behavioral and spiking data was performed using custom MATLAB code (Mathworks, MA).

### Task

#### Navigate-To-Sample (NTS)

Each trial begins with the animal fixating a central spot of size 0.5 degrees of visual angle (dva). Next, an image sequence is presented above the fixation point. The sequence contains 6 equidistant landmark images (inter-landmark distance of 6 dva), denoted *I*_*1*_ to *I*_*6*_. The sequence is shifted horizontally such that a randomly chosen image from the sequence (*I*_*i*_) appears directly above the fixation point. We refer to this landmark as the initial landmark. Next, we present a different randomly selected image from the sequence (*I*_*j*_) directly below the fixation point, which we refer to as the terminal landmark. The initial and terminal landmarks stay on the screen for a variable time (400-1400 ms, uniform hazard). Afterward, a change in the color of the fixation point serves as the Go cue instructing the animals to deflect a 1D joystick to move the entire sequence above the fixation leftward or rightward at a constant speed (10 dva/s) and stop when the image right above the fixation point matches the terminal landmark image below the fixation point (see Fig. S1a and Supplementary video 1). Throughout the trial, only images that were within the width of the monitor were visible. Trials are separated by an inter-trial interval (ITI; 500-1000 ms, uniform hazard). In essence, animals must produce a 1D vector *v*_*p*_ that matches the vector extending from the initial landmark to the terminal landmark, denoted *v*_*a*_. Since the movement speed is constant, these vectors can be expressed as signed numbers whose magnitude corresponds to the temporal distance between images and whose sign represents direction. We designated rightward and leftward pointing vectors as positive and negative, respectively.

#### Mental-Navigation (MNAV)

We used NTS to help the animals learn the basic task contingencies, inter-landmark distance, image sequence, and joystick speed. Next, we started the training on the main Mental-Navigation (MNAV) task, which is identical to NTS except for the following two critical differences. First, during the initial presentation of landmarks (before the Go cue), all the landmarks on the sequence except the one right above the fixation are invisible. Second, between joystick onset and offset, all landmarks, including the one above the fixation, are made invisible. After the joystick offset, the landmark closest to the fixation point is presented (Fig 1a and Supplementary Video 2). Animals receive reward if the relative error defined as |*v*_*p*_-*v*_*a*_|/*v*_*a*_ is smaller than a criterion value of .08*|*v*_*a*_|. If not, the animals are given a second and last chance to produce a corrective vector. The second attempt furnishes 1/4 of the original reward if the relative error is smaller than the criterion. When rewarded, reward decreased linearly with relative error. When animals aborted the trial by deflecting the joystick before the go cue, a timeout of 5 sec was added to the ITI. This was done to discourage the animals from purposely aborting long-vector trials.

### Electrophysiology

We located EC based on stereotaxic coordinates and structural MRI scans of the two animals ^70^. To target EC reliably, we used a grid system inside the recording chamber. We registered grid holes relative to the brain using an MRI scan in which the holes were filled with an MRI contrast agent (5mg/ml Gadolinium + 10mg/ml agar). We used the registered grid system together with readings of anatomical landmarks along the penetration path to target EC accurately. We recorded extracellularly across 32 sessions (A:17, M:15). Recorded signals were amplified, bandpass filtered, sampled at 30 kHz, and saved using the OpenEphys data acquisition system (OpenEphys Inc., Lisbon, Portugal). Using Kilosort 2.0 software ^71^, we isolated 1478 single- and multi-units (A:614, M:864).

### Analysis of neural data

We focused our neural data analysis on the subset of task-modulated neurons defined as those whose firing rate either exhibited periodicity during navigation or was modulated by temporal distance during joystick onset or offset (see below for the corresponding analyses). To plot firing rates, we smoothed spike counts in 1-ms bins using a Gaussian kernel with a standard deviation of 100 ms. Because of variability in the produced vectors, trials associated with the same condition (i.e., same direction and distance) had variable lengths. To compute trail-averaged firing rates for each condition, we used 40 ms bins for the median produced interval, and appropriately stretched or compressed bins for shorter and longer trials ^65^.

### Periodicity

We computed a periodicity index (PI) for each EC neuron using a similar procedure used for computing gridness score during spatial navigation tasks ^36,55^. (1) We pooled firing rates for trials requiring mental navigation over at least three landmarks to ensure that trials were long enough to compute periodicity. (2) We truncated trials at 500 ms before the joystick offset to ensure that our estimate of periodicity was not biased by the associated large anticipatory response (see Fig 2a). (3) We detrended firing rates using linear regression so that ramping activity would not mask the presence of periodicity (Fig S8). (4) We computed an average autocorrelogram (ACG) for each neuron by averaging the single-trial autocorrelation function of firing rates at lags between 0 and 2400 ms. (5) To detect periodicity, we computed the correlation between the ACG and shifted ACG for varying lags ranging from 0 to 1300 ms. (7) We defined PI at each lag as the difference between ACG for that lag and ACG for half that lag. This procedure is analogous to how gridness scores are computed, with the difference that instead of a two-dimensional spatial ACG, a one-dimensional temporal ACG is used.

To evaluate the significance of PI for each neuron, we also created a null distribution for PI using surrogate data generated from a zero-mean Gaussian Process (GP) with a squared exponential kernel (maximum variance =1; length constant = 100 ms). To match the smoothness of GP to our smoothed firing rate, the length constant parameter of the squared exponential kernel was equal to the width of the Gaussian smoothing kernel (100ms) used for smoothing the firing rates of EC neurons. We then passed the GP surrogate data through a non-homogenous Poisson process and smoothed the resulting spike train to obtain our surrogate GP null data. We repeated this process 1000 times to obtain a distribution. A neuron was classified as periodic if its PI at any lag was higher than 2 standard deviations from PI obtained from surrogate data. For neurons with significant PI, the preferred period was the lag at which the PI was the maximum.

### Temporal distance modulation

For each neuron, we also quantified the degree to which firing rates at joystick onset and offset were modulated by the produced temporal distance. For each neuron, we sorted the trials into 15 bins according to produced vector length (*v*_*p*_), and computed the regression coefficient relating those lengths to the corresponding firing rates, both at joystick onset and offset. A neuron was considered to exhibit significant temporal distance modulation if the regression slope at either onset or offset were significant (F-statistic at 95% confidence).

### Quantification of the phase of local bumps of activity

To find the phase of localized bumps of activity with respect to joystick offset, we detected the phase of local maximum within a 1-second window before joystick offset. We bootstrapped over 100 repeats of subsampled trials and calculated the distribution of the mean phase. We created a null dataset by circularly shifting each trial’s spike train by a random length and repeating the phase detection steps.

To find the phase of the second-last peak, we detected the phase of the local maximum in the 1-second window before the first peak relative to the joystick offset. Similarly, the phase of the third-last peak was the phase of the local maximum in the 1-second window before the second peak relative to the joystick offset.

We computed the mode of the absolute phase distribution for each periodic neuron whose phase distribution was different from its shuffled control (KS test, p<<<.001).

### Targeted dimensionality reduction

We used a regression analysis across the population of neurons to identify the dimension in which firing rates encode temporal distance ^39^. We first centered the responses of each neuron by subtracting its mean response across the window of interest (500ms before joystick onset and offset). We then computed the regression line relating the firing rates to the temporal distance:

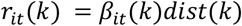

Here, *disk*(*k*) is the ordinal distance on trial k (dist: 1 to 5) and *r*_*it*_(*k*) is the centered firing rate for trial k at time t and for neuron i. For N neurons we then had a Nxt matrix of regression coefficients. We took the norm of this matrix along the dimension of time to find the time point where the coefficients were maximum. The vector of regression coefficients is considered the optimal axis for coding ordinal distance. Next, we projected the matrix of held-out trials for the same neurons onto the optimal distance axis (Fig3c,d).

Linear encoding of distance was quantified by measuring the variance accounted for by a linear model (R^square^) with a dummy independent variable, dist = [1,2,3,4,5]. To compare the encoding of the ordinal distance across the two epochs - joystick onset and offset, we created a bootstrapped distribution (50 resamples) of distance projections averaged over a 100 ms window before joystick onset and offset.

### Continuous attractor model

We tackled the circuit-level modeling of our task in two steps. First, we examined the conditions under which grid cell activity could serve as internal landmarks. To do so, we constructed a model with multiple GC modules with different periodicities and phases, and a hypothetical landmark neuron (LM). LM received both external input (*I*_*ext*_) from visual stimuli and internal input (*I*_*int*_) from GCs. The synaptic weights from GC to LM were subjected to plasticity. Learning proceeded in two stages, first with both *I*_*ext*_ and *I*_*int*_, mimicking conditions in NTS, and later with *I*_*int*_ only, mimicking conditions in MNAV. Initially, synaptic weights were sampled randomly from a normal distribution. Throughout learning, synapses were updated at every time step using Oja’s rule. To ensure learning was stable, we (1) used a sufficiently small learning rate (i.e., smaller than 1e-7) and (2) normalized the weights for each module such that they were always centered at zero. This learning scheme selectively strengthened inputs from the subset of GCs whose periodicity and phase match external input (Fig. 4, S7).

Having established that learned *I*_*int*_ could emulate *I*_*ext*_, we next constructed a continuous attractor network of GCs adapted from Burak and Fiete^20^ (code at: https://fietelab.mit.edu/code/) to compare the effect of attractor dynamics in path integration versus mental navigation. The model GCs have difference-of-Gaussian connectivity kernels, with centers shifted clockwise or counterclockwise, and performs path integration when driven by matching velocity inputs (e.g. left-shifted cells receive leftward velocity inputs). We first characterized the path integration behavior of this model in the presence of variable velocity input. To do so, we simulated the network when the velocity input on each trial was perturbed by Gaussian noise, and quantified the time it takes for the network to reach different distances measured in terms of network phase (Fig. 4c, black).

Next, we did the same for mental navigation behavior by providing the additional *I*_*int*_ from the subset of GCs whose periodicity and phase match *I*_*ext*_ (Fig. 4c, red). Finally, we compared the overall variability of the model in the two contexts; i.e. path integration versus mental navigation (Fig. 4d).

### Bayesian model of behavior

We fit two Bayesian observer models to behavioral data (Fig S8a). Both models combine the likelihood function associated with a noisy measurement with the experimentally-imposed prior distribution and use the posterior mean, *t*_*e*_ (i.e., Bayes least squares estimation) to estimate the temporal distance ^72^. Both models also assume that the navigation epoch introduces additional variability. The variability in the path integration model is assumed to follow scalar property (Fig S8a) ^51,52^ i.e., standard deviation increases linearly with base interval:

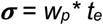

In contrast, the mental navigation model divides longer temporal intervals to multiples of the base interval (*t*_*o*_), which in our experiment is 0.65 sec. In this case, by variance sum law, the total variance ***σ***_*2*_ is:

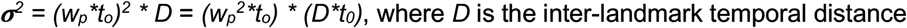

Accordingly, the standard deviation grows as the square root of the total interval:

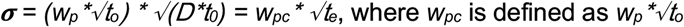

We first used surrogate data generated by each model to verify that our fitting procedure could correctly identify the generative model (Fig S8 b,c). Next, we compare the model fits to behavior. The models used to fit the behavior were augmented to include an offset term, *b*, to account for the overall bias in produced temporal intervals. We used maximum likelihood estimation (MLE) to fit the models to behavior on each session, and used the predicted bias and variance to compare the models.

## Supplementary figures

**FigS1.**
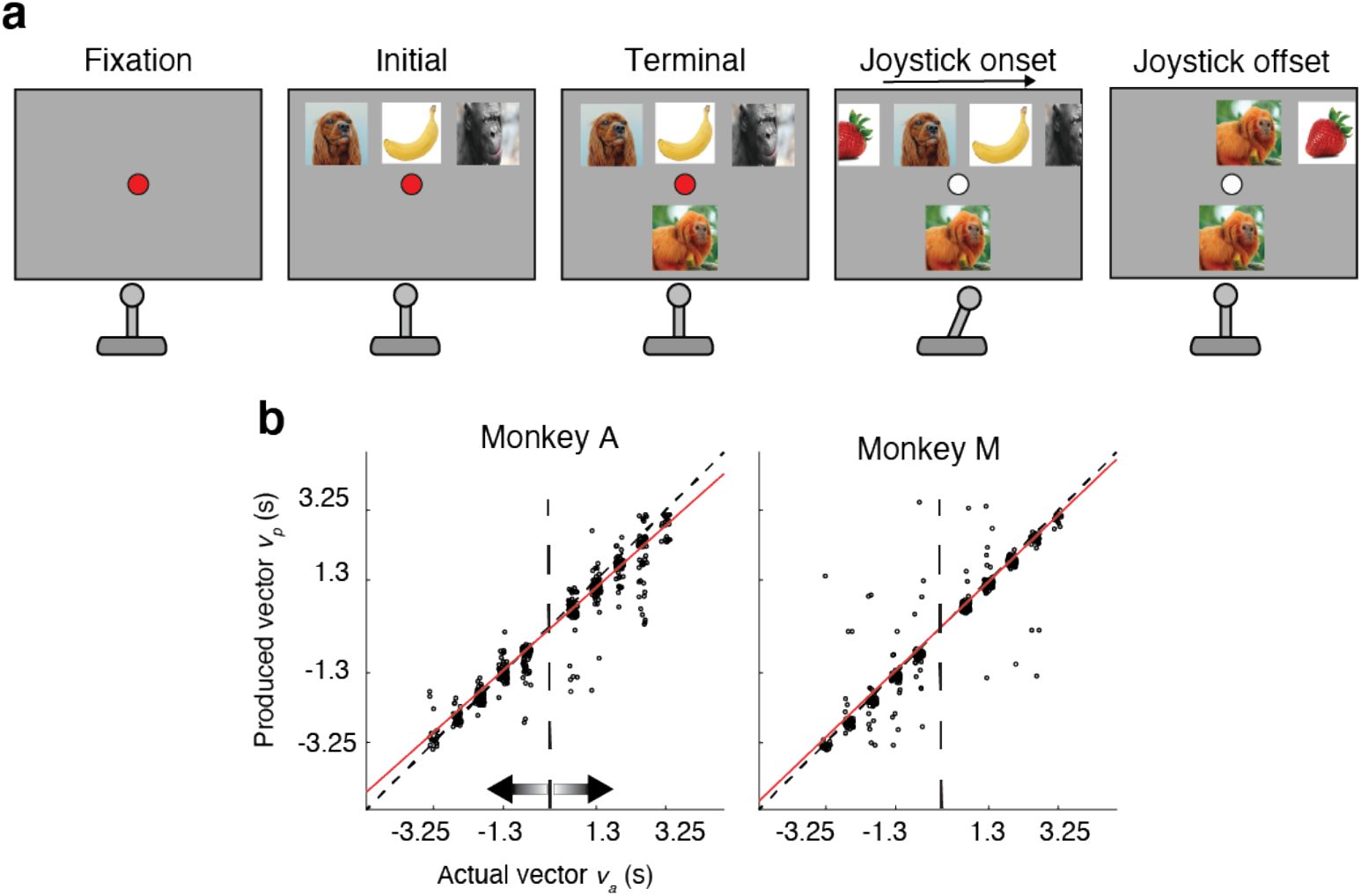
Navigate-to-sample task and behavior. **a**, Sequence of events during a trial of the navigate-to-sample (NTS) task. The sequence is identical to the MNAV task (Fig. 1a), except that during joystick deflection, all images and their movement are visible. **b**, Behavior in a representative session quantified in the same manner as the MNAV task (Fig. 1b).

**FigS2.**
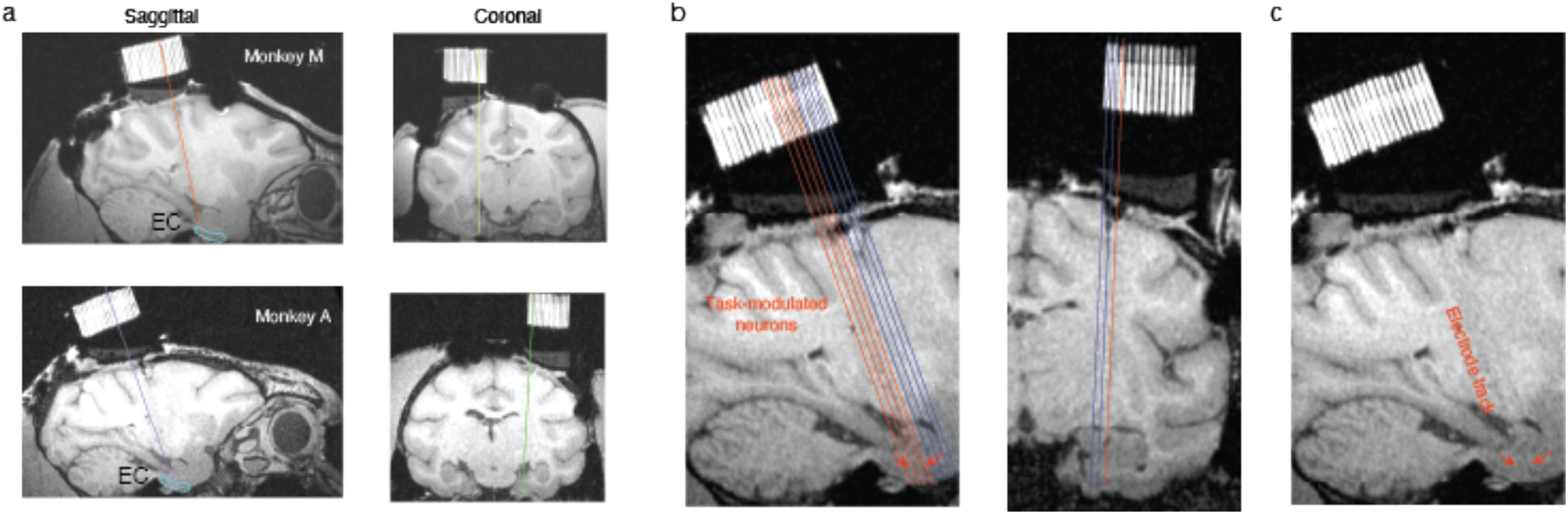
Recording site. **a**, Structural scan of a monkey M (top) and monkey A (bottom) together with the recording grid across sagittal and coronal anatomical planes. **b**, Tracks through the grid (red: tracks with task-modulated neurons). Anatomical coordinates: 9.5-10.5 mm from the midline, 14-16 mm anterior to *ear bar zero* (EBZ), and centered on EBZ along the vertical axis. **c**, Post-recording scan showing electrode tracks and recording sites targeting periodic neurons in EC.

**FigS3.**
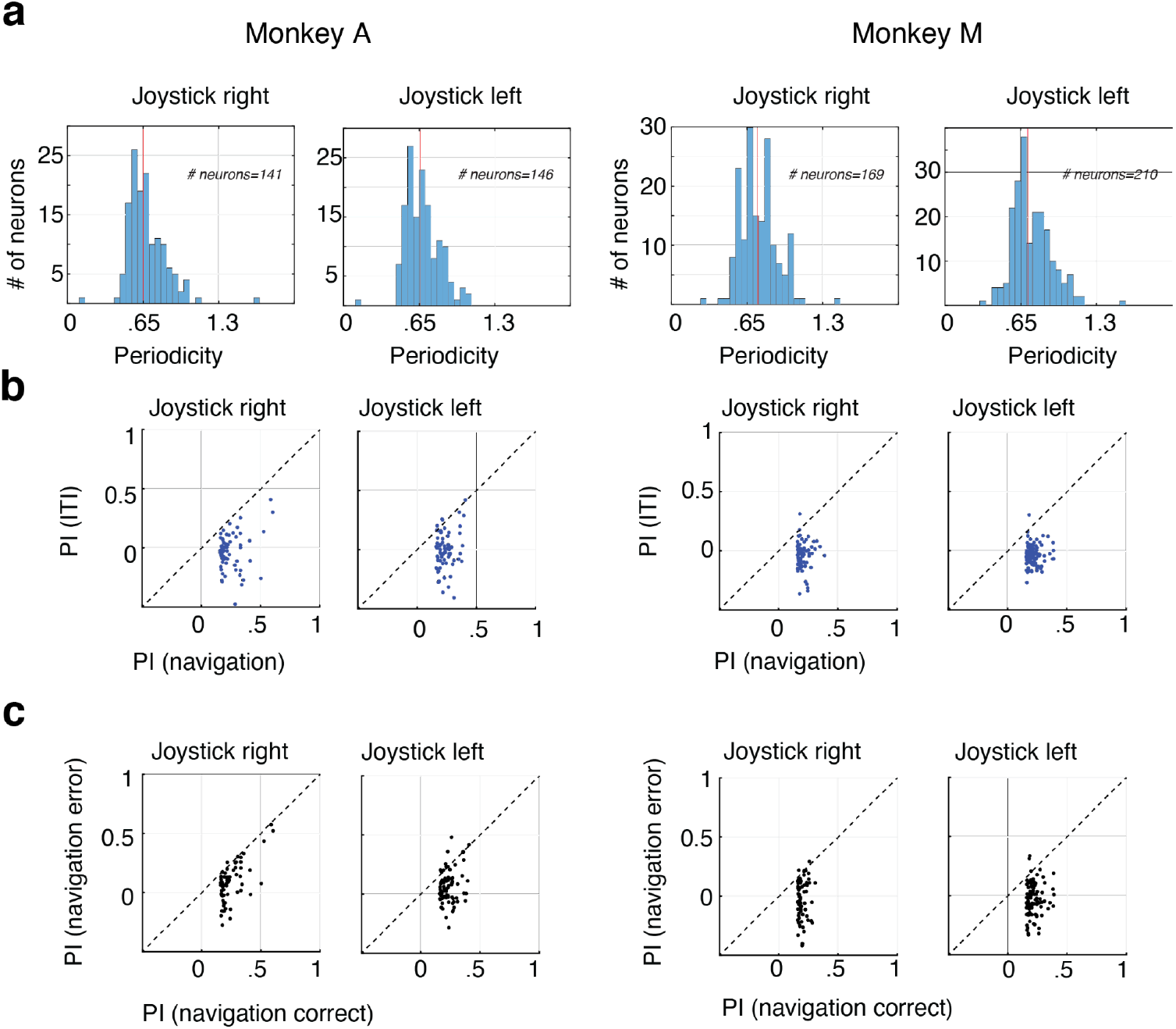
Periodicity across sessions for different directions of navigation, epochs, and trial types. **a**, Distribution of periodicity of all periodic neurons for two animals and for two directions of navigation. **b**, Comparison of periodicity index at 0.65 sec during the inter-trial interval (ITI) versus the navigation epoch. One-tailed paired t-test, monkey A, right: t(67)=14.07, p<<.0001; A, left : t(75)=14.02, p<<.0001; monkey M, right: t(73)=16.93, p<<.0001; monkey M, left: t(103)=26.56, p<<.0001. **c**, Comparison of periodicity index at 0.65 sec during error versus successful trials in the navigation epoch. One-tailed paired t-test, monkey A, right: t(67)=10.58, p<<.0001; A, left : t(75)=11.02, p<<.0001; monkey M, right: t(73)=10.11, p<<.0001; monkey M, left: t(103)=17.73, p<<.0001.

**FigS4.**
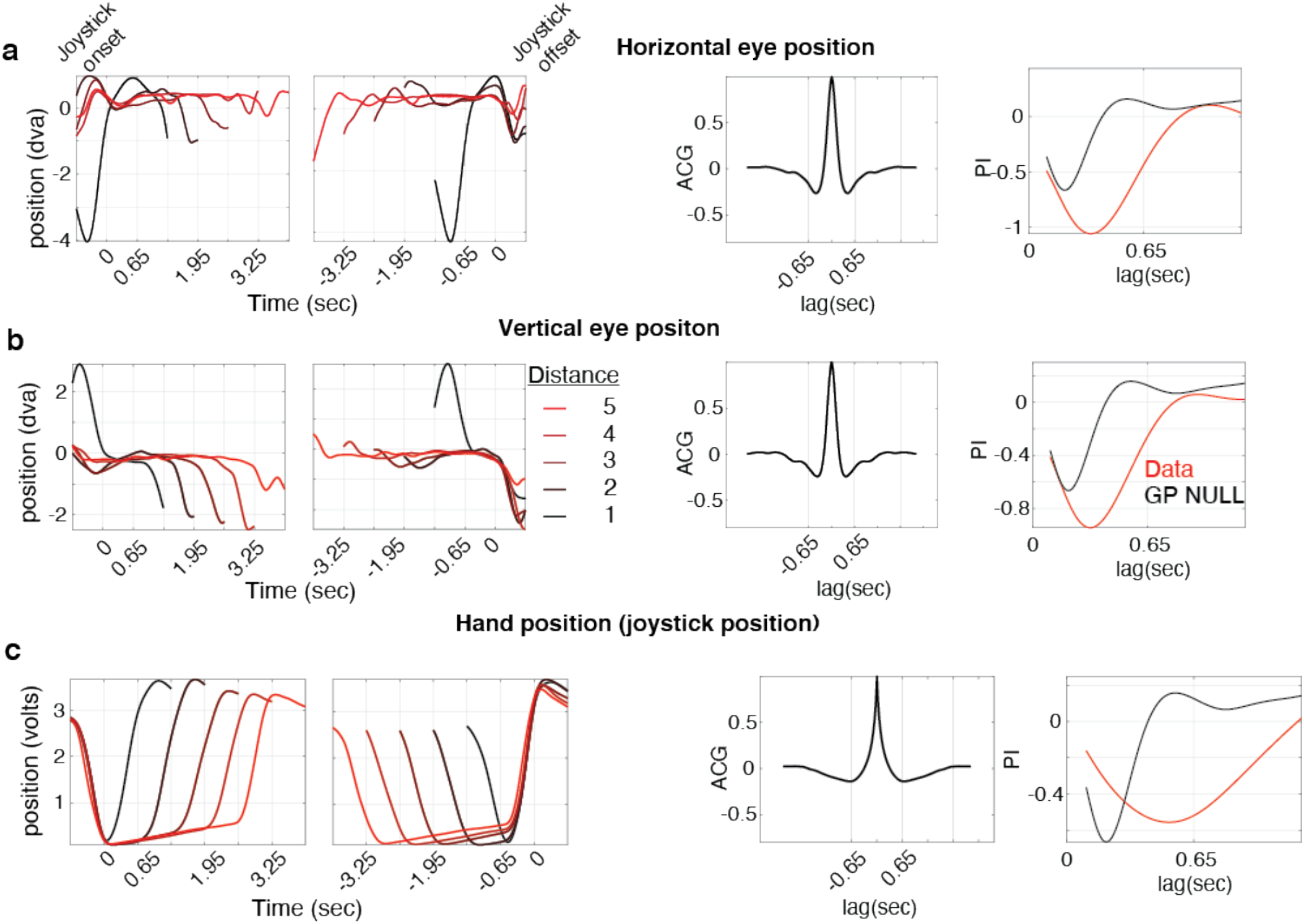
Eye and hand position time course dynamics during the navigation epoch. **a**, Left. Time course of eye position between joystick onset and offset averaged across trials for horizontal eye position disaggregated for five different temporal distances (shades of red). Middle. Average autocorrelogram of the horizontal eye position time course. Right. Periodicity indices (PI) at different time lags for horizontal eye position together with the significance threshold (black) at 2 standard deviations above chance-level PI computed on Gaussian Process surrogate data (n=100 bootstraps, average PI at 650ms = −0.25 is smaller than chance (one-tailed t-test (99) = −67.8, p<<.0001). **b**, Same as **(a)** for vertical eye position (n=100 bootstraps, average PI at 650ms = −0.44 is smaller than chance (one-tailed t-test (99) = −81.6, p<<.0001). **c**, Same as **(a)** for hand position measured with the voltage readout of joystick position (n=100 bootstraps, average PI at 650ms = −0.53 is smaller than chance (one-tailed t-test (99) = -138.1, p<<.0001).

**FigS5.**
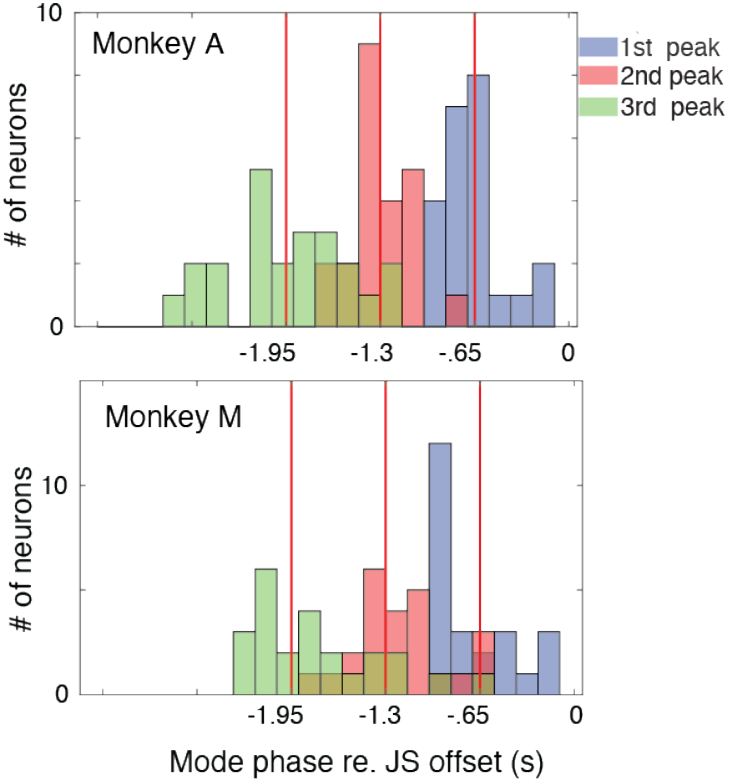
Relationship of peak phase to behavior for two animals plotted separately. Same as Fig. 3F for the two monkeys separately. Distribution of the first (blue), second (red), and third (green) peak phase before joystick offset across neurons for monkey A (top) and monkey M (bottom).

**FigS6.**
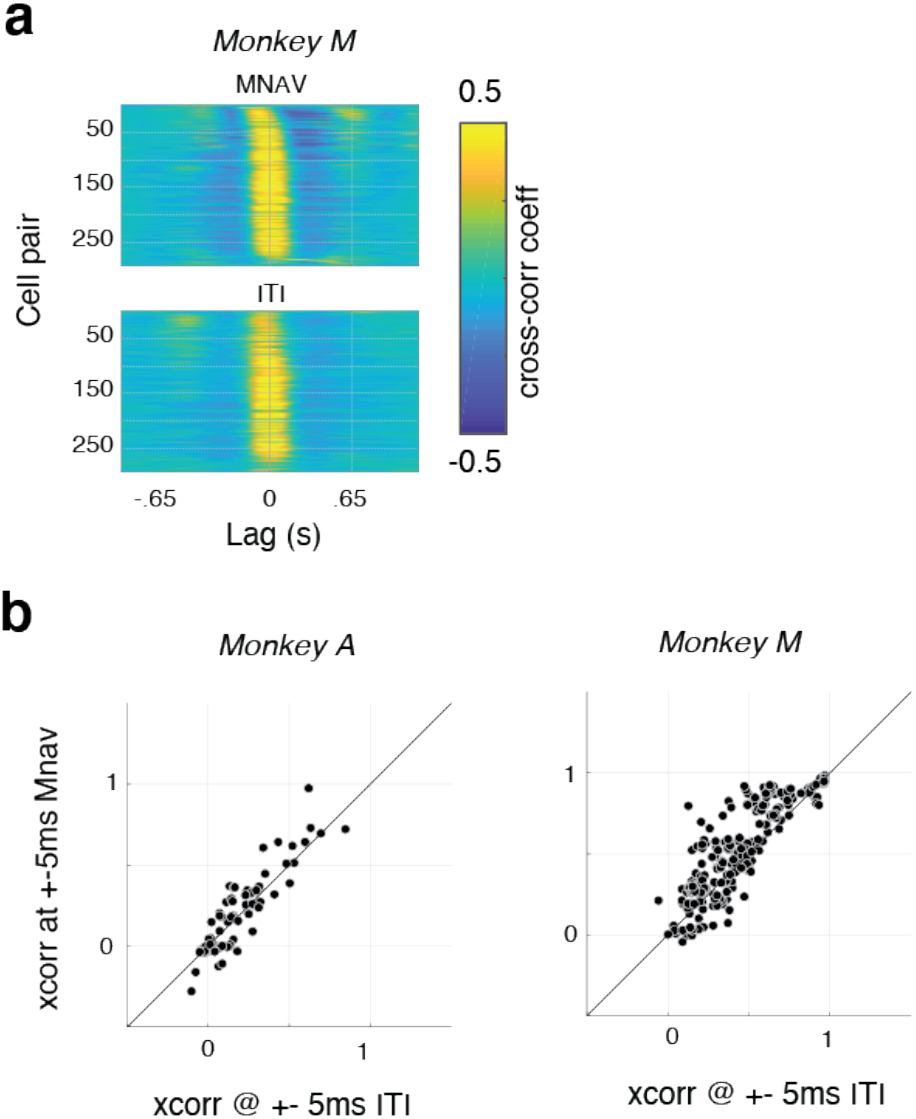
Cross-correlation structure of periodic neurons across epochs in the second animal. **a**, Cross-correlation structure of 25 simultaneously recorded periodic neurons with the highest periodicity metric (292 pairs) rank ordered based on activity during the navigation epoch (top: MNAV) and plotted with the same order for the inter-trial interval (bottom: ITI) for monkey M. **b**, Scatter plot of firing rate cross-correlation averaged over a lag window of +-5ms in the two different epochs for monkey A (left) and monkey M (right).

**FigS7.**
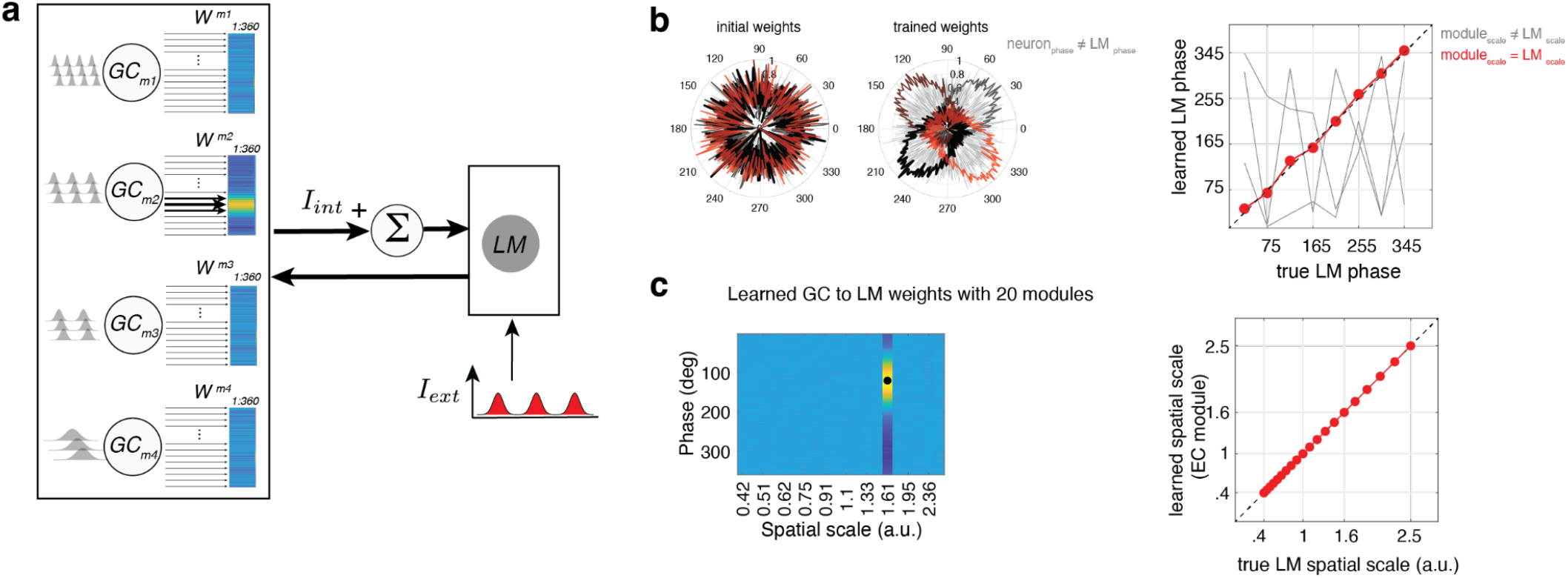
Model identifiability for continuous attractor network model. **a**, Schematics of the model similar to Fig. 4a. **b**, (Left, middle) Learning of GC to LM connections in the presence of *I*_*ext*_ with a specific periodicity and 4 different phases (i.e., 4 different model simulations). Synaptic weights from all neurons in all GC modules (m1, m2, m3, m4) are initially random (left). Through Hebbian learning synaptic weights of those GC cells whose periodicity and phase match *I*_*ext*_ (middle). (Right) 8 simulations of the model, each with the landmark input at a chosen phase. Learned landmark phase, computed as the phase of the GC neuron with maximum weight to the LM neuron, plotted as a function of the true landmark phase for the appropriate module (red) and all other modules (gray). **c, (**Left) In the presence of *I*_*ext*_ with a specific periodicity and phase (black circle), a new instantiation of the model with 20 modules learns the right phase and the right periodicity. (Right) Across 20 simulations of the model, each with the landmark input at a chosen periodicity learned landmark spatial scale (module) plotted as a function of true landmark spatial scale.

**FigS8.**
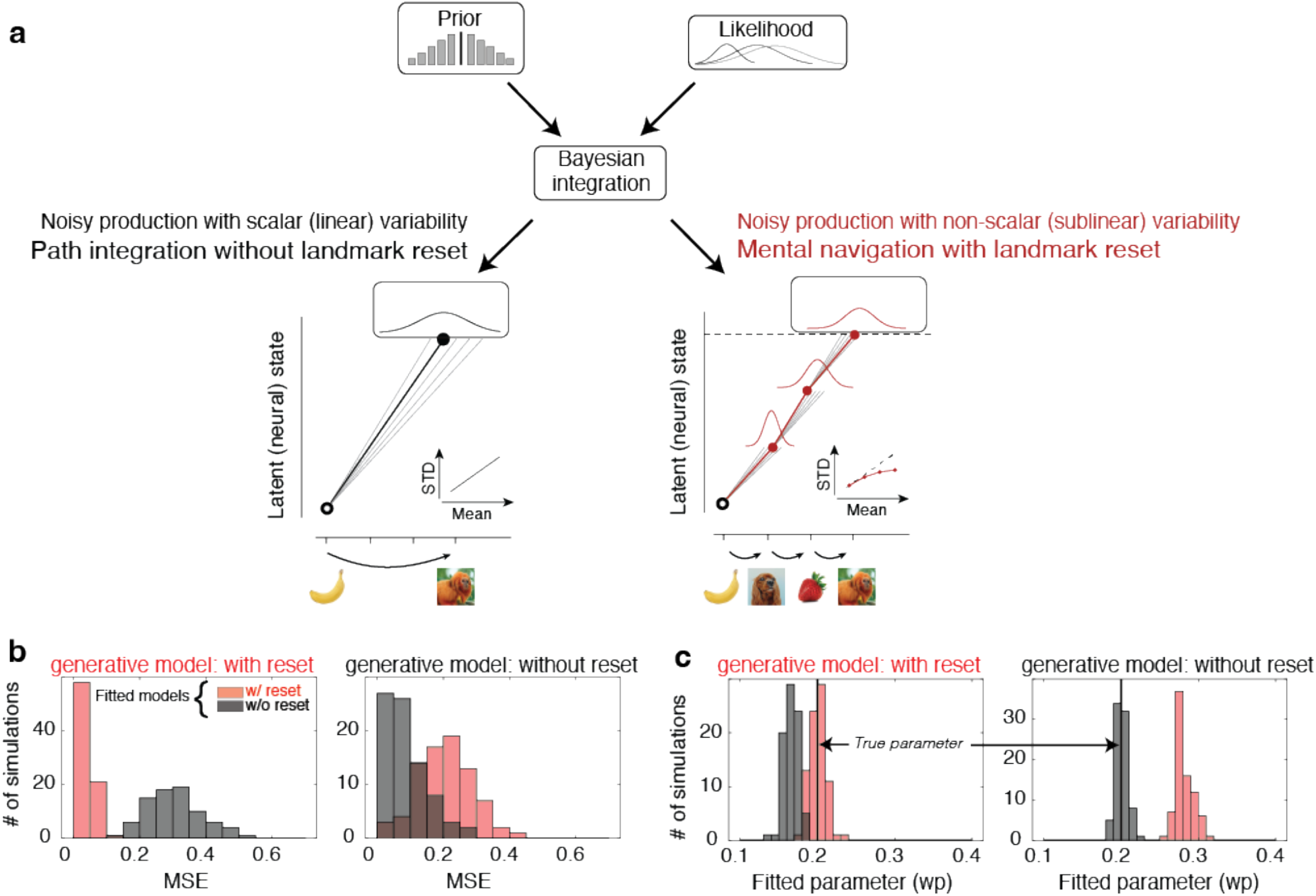
Description and identifiability of the Bayesian observer models. **a**, Schematic illustration of the Bayesian observer model that combines the prior (top left) with noisy estimates of temporal distance. 3 likelihood functions displayed associated with three different noisy estimates (top right). The output of the Bayesian integration, the posterior mean undergoes noisy production during the mental navigation epoch. Two models of noise accumulation: left. Path integration without landmark reset where standard deviation increases linearly with temporal distance. Right: Mental navigation with landmark reset where standard deviation increases sublinearly with temporal distance (i.e., variance increases linearly with temporal distance). **b**, Distribution of mean squared error (MSE) between ground truth data generated from a chosen generative model and data generated from the two models fitted to the ground truth data. Left: ground truth data generated from a model with reset (blue). Right: ground truth data generated from a model without reset (red). **c**, Distribution of fitted parameter values (Weber fraction, w_p_) for the two fitted models when data was generated by the model with reset (left) and when the data was generated by the model without reset (right).

## Acknowledgments

SN is supported by NSERC PDF-516867-2018, FRQNT B3X-258512-2018. IF is supported by NIH (NIMH-MH129046). MJ is supported by NIH (NIMH-MH129046), Paul and Lilah Newton Brain Science Award, and the McGovern Institute.

## Notes

### Competing Interest Statement

The authors have declared no competing interest.

